# Autofluorescence lifetime imaging resolves cell heterogeneity within peripheral blood mononuclear cells

**DOI:** 10.64898/2026.03.06.710224

**Authors:** Jeremiah Riendeau, Lucia Hockerman, Elizabeth Maly, Kayvan Samimi, Melissa Skala

## Abstract

**Significance:** Standard methods to characterize peripheral blood mononuclear cells (PBMCs) are often destructive, lack metabolic information, or do not provide single-cell resolution. Label-free tools that non-destructively measure single-cell metabolism within PBMCs can provide new layers of information to characterize disease state and cell therapy potential.

**Aim:** Determine whether non-destructive fluorescence lifetime imaging microscopy (FLIM) of endogenous metabolic co-factors NAD(P)H and FAD, or optical metabolic imaging (OMI), can identify immune cell subsets and activation state within heterogeneous PBMC cultures.

**Approach:** OMI measured single-cell metabolism of PBMCs from 3 different human donors in the quiescent or activated (phorbol 12-myristate 13-acetate and ionomycin) state. Fluorescent antibodies were used as ground truth labels for single-cell classifiers of immune cell subtypes.

**Results:** OMI identified quiescent vs. activated PBMCs with 93% accuracy at only 2 hours post-stimulation, identified monocytes within quiescent and activated PBMCs with 96% and 88% accuracy, respectively, and identified NK cells within quiescent and activated PBMCs with 74% accuracy.

**Conclusion:** OMI identifies activation state and immune cell subpopulations within PBMCs, enabling single-cell and label-free measurements of metabolic heterogeneity within complex PBMC samples. Therefore, OMI could enhance PBMC immunophenotyping for diagnostic and therapeutic applications.

**Statement of Discovery:** We demonstrate that autofluorescence lifetime imaging can resolve functional and phenotypic metabolic subpopulations within a mixed culture of immune cells from human blood. This provides a new technique to characterize metabolic activity within immune cells from the peripheral blood of patients, which could improve disease diagnostics and the production of cell therapies.

## Introduction

Peripheral blood mononuclear cells (PBMCs) are a diverse cohort of immune cells comprised of lymphocytes (T cells, B cells, and NK cells) and myeloid cells (monocytes) [1]. Due to the high degree of immunologic heterogeneity, PBMCs are used in many applications from modeling the immune system to monitoring disease progression [2,3]. Additionally, PBMCs are often used as a starting material for cell therapies [4,5]. PBMCs also provide an easily accessible immune cell population that is rapidly isolated from whole blood through simple density gradient centrifugation. PBMC samples are most often assessed using flow cytometry, which provides the proportion of lymphocytes and monocytes within the PBMC sample via forward-scattering and side-scattering [1,6]. Flow cytometry of fluorescently labeled surface markers can further identify populations of T cells, B cells, NK cells, and monocytes while providing functional measurements such as activation state. However, fluorescent labels can interfere with biological processes and require lengthy staining procedures that can alter cell features [7–10]. Additionally, standard flow cytometry does not routinely assess cell metabolism, yet metabolic state reveals unique cell subpopulations and cell functional states compared to surface markers alone [11–14].

Quiescent PBMCs are not only comprised of functionally diverse cell subsets, but also metabolically distinct cell subsets. Quiescent T cells generate the majority of their adenosine triphosphate (ATP) through oxidative phosphorylation (OXPHOS) via fatty acid oxidation (FAO) [16–17]. Quiescent lymphocytes (majority T cells) treated with oligomycin were unable to increase glycolysis, further supporting that T cells rely almost exclusively on FAO/OXPHOS for energy production with little room for metabolic flexibility [18]. Like quiescent T cells, quiescent B cells take up little glucose, maintain few, small mitochondria and exhibit an extremely low metabolic rate, making both quiescent T and B cells metabolically quiet [19,20]. As an innate immune cell, monocytes are primed for rapid activation and therefore maintain a higher metabolic rate in the quiescent state compared to quiescent lymphocytes [18]. While quiescent monocytes do rely on FAO/OXPHOS for the majority of ATP production, glycolysis and glutaminolysis remain important secondary energy sources that allow monocytes to rapidly switch to glycolysis with activation [18]. NK cells are classically described as innate immune cells like monocytes, though are derived from the same lymphocyte lineage as B and T cells [21]. As such, NK cells maintain an intermediate level of metabolic activity with higher OXPHOS reliance than monocytes though less than T and B cells, while also using glucose as an important secondary energy source [22–24]. With activation, NK cells show increased rates of both glycolysis and OXPHOS with high production of lactate [22–24]. Post-activation, cell metabolism is heterogeneous with glycolysis required for inflammatory function in several immune cells including CD8 T cells, monocytes, and NK cells [24–26], while OXPHOS is required for B cell activation, immunosuppressive functions (e.g., regulatory T cells), and for memory formation [19, 27, 28].

Immune cell metabolism is emerging as a key feature of immune cell behavior and disease state because metabolism more closely aligns with temporal changes in immune cell function compared to surface markers that exhibit delayed expression [29,30]. PBMC metabolic states have already demonstrated success in predicting heart failure severity [31], cognitive decline [32,33], cancer treatment response [34,35], and severity of inflammatory conditions like sepsis and systemic lupus erythematosus (SLE) [36–38]. PBMC metabolism has also been correlated with fitness level and physical frailty, revealing PBMCs as an important marker of whole-body metabolic fitness [39–42]. Metabolism has also identified heterogeneity within subsets of immune cells that cannot be defined by surface marker expression alone, for example by identifying metabolic markers of more successful chimeric antigen receptor (CAR) T cell therapies [13] and identifying the role of glutamine-dependent T and B cells that fuel inflammation in SLE, offering promising therapeutic targets [43,44]. Therefore, tools that can measure single-cell metabolism within PBMCs could provide valuable information about immune cell function and heterogeneity beyond traditional surface marker expression.

Fluorescence intensity and lifetime imaging of the metabolic cofactors, nicotinamide adenine dinucleotide (phosphate) (NAD(P)H) and flavin adenine dinucleotide (FAD), or optical metabolic imaging (OMI), provides single-cell metabolic imaging of individual cells [45–47]. NAD(P)H exists in a free and protein-bound state that exhibits a short and long fluorescence lifetime, respectively [48,49]. Conversely, the free and protein-bound states of FAD have a long and short fluorescence lifetime, respectively [50]. These lifetimes are sensitive to the molecular microenvironment (e.g., pH, viscosity, temperature, ionic strength) [51–53] while the protein-bound lifetime also varies by binding partner [54–56]. Additionally, the fluorescence intensities of NAD(P)H and FAD are used to calculate the optical redox ratio (ORR) [57–59]. While the ORR has many definitions, here the ORR is defined as the intensities of NAD(P)H / [NAD(P)H + FAD]. OMI relies on intrinsic sources of contrast, and this label-free imaging technique is attractive because no sample manipulation is required. Therefore, the same cells can be repeatedly imaged within intact cultures over time before performing endpoint analyses, or cells can be imaged before use in cell manufacturing or cell therapy.

OMI has already successfully classified quiescence and activation in monocultures of primary human neutrophils [60], T cells [61–63], B cells [64], and NK cells [64], and classified CD4+ and CD8+ T cells within bulk CD3+ T cell cultures [63]. Autofluorescence signals have also revealed differences between several immune populations present in peripheral blood including lymphocytes, monocytes, and several granulocytes [65,66]. While previous studies explored autofluorescence characteristics of immune cells in monoculture or quiescent systems only, this paper is the first to show that OMI can classify activation state and immune cell subpopulations within PBMCs from primary human donors. This could provide a new tool to measure single-cell metabolism within PBMCs and thereby provide label-free metabolic information to better monitor diseases characterized by immune activation (e.g., sepsis, SLE, rheumatoid arthritis) and screen PBMCs for cell therapy (e.g., verifying metabolic fitness of starting cells and products).

## Methods

### 1.1 Cell Preparation

PBMCs were isolated from the peripheral blood of healthy adult donors under approval by the University of Wisconsin-Madison Institutional Review Board. All donors gave written informed consent in accordance with the Declaration of Helsinki. After obtaining written informed consent from the donors, 10 mL whole blood was drawn using a sterile syringe into a vacutainer tube coated with sodium heparin (158 United States Pharmacopeia units). Whole blood was mixed with an equal volume of Dubelco’s phosphate buffered saline (DPBS) + 2% fetal bovine serum (FBS) and layered on top of 15 mL Lymphoprep density gradient (Stemcell Technologies) in a 50 mL SepMate tube (Stemcell Technologies). Tubes were centrifuged for 10 minutes at 1200xg with no brake. The yellow plasma layer was aspirated and the cloudy PBMC layer was poured off into a new tube and washed with DPBS + 2% FBS, centrifuged for 5 minutes at 800xg, with the brake on. The PBMC pellet was resuspended at 2×10^6^ PBMCs/mL of media (RPMI 1640 + 2mM glutamine + 10% FBS + 1% penicillin/streptomycin).

PBMCs were plated at 500k PBMCs in 200 μL media into wells of a poly-D-lysine coated glass 96 well plate without activators (quiescent) or with 40 nM phorbol 12-myristate 13-acetate (PMA) and 1 μM ionomycin (activated). After plating, each well of mixed PBMCs was incubated for 30 minutes with 1.0 μL of one fluorescent antibody per well (PerCP-Cyanine5.5-αCD4, PerCP-Cyanine5.5-αCD8, PerCP-αCD14, Star Bright Blue 700-αCD19, Brilliant Blue 700-αCD56) [**Fig. 1(a)-(b)**] [**Supp Table 1**] to identify a single cell population within the mixed culture [**Fig. 1(c)**]. Excess fluorescent antibody was removed by centrifuging cells for 30 seconds at 800xg and discarding supernatant. Cells were washed 3x with PBS + 2% FBS and finally resuspended in 200 μL media. After 2 hours from the initial plating, quiescent and activated PBMCs were imaged using OMI.

**Figure 1.**
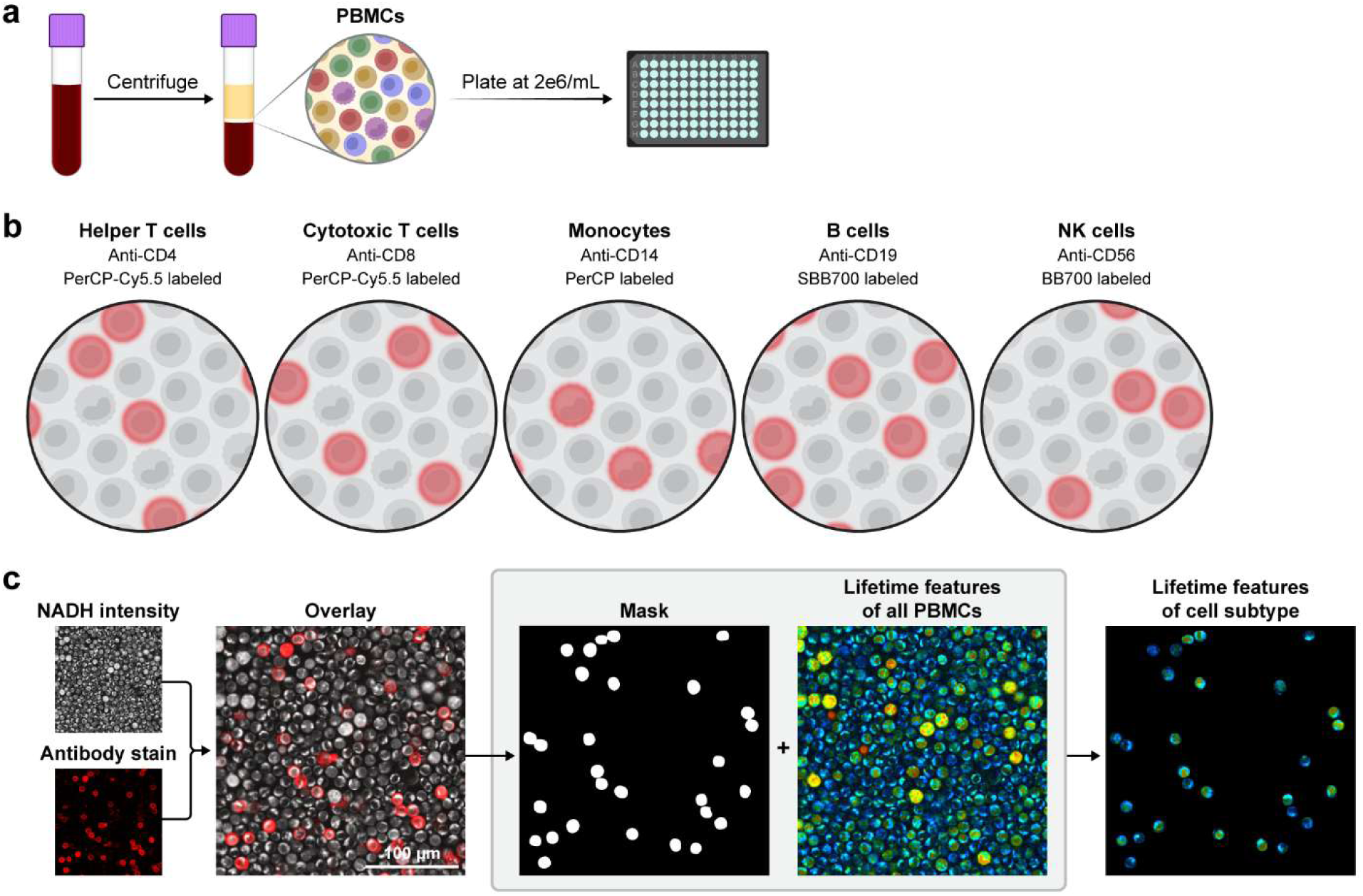
PBMC staining and image processing workflow. (a) PBMCs are isolated via density centrifugation of whole peripheral blood and plated in wells of a 96 well plate. (b) One fluorescent antibody is added per well to identify single cell types within the mixed culture. The fluorophores PerCP, PerCP-Cy5.5, star bright blue 700 (SBB700), and brilliant blue 700 (BB700) were used as fluorescent reporters. (c) Single cell masks are generated from the stain intensity and NAD(P)H intensity images to extract OMI features of the specified cells.

### 1.2 Optical Metabolic Imaging

PBMCs were kept in a stage top incubator (Tokai Hit) at 37°C and 5% CO_2_ for the duration of imaging. Cells were imaged on a custom-built two-photon microscope (Bruker) using Prairie View Software (Bruker) with a 40x (NA = 1.15) water-immersion objective (Plan Apo, Nikon) with a 1.5x digital scan zoom. An Insight DS+ tunable femtosecond-pulsed laser (Spectra-Physics Inc.) with 80 MHz pulse repetition rate was tuned to 750 nm to excite NAD(P)H and the fluorescently labeled antibodies with an average power of 6.0 mW at the sample, while 890 nm was used to excite FAD with an average power of 15.0 mW at the sample. The laser power was maintained at a consistent value within each experiment. Fluorescence emission was detected using a H7422PA-40 GaAsP photomultiplier tube (Hamamatsu) and isolated using a 466/40nm bandpass filter for NAD(P)H fluorescence, a 520/40 nm bandpass filter for FAD, and a 590 long pass filter to collect fluorescence of labeled antibodies. Fluorescence decays were captured using a Time Tagger Ultra (Swabian Instruments GmbH). Photon count rates were maintained at 1.5×10^5^ photons/second to ensure reliable lifetime fits.

A pixel dwell time of 4.8 μs and an integration time of 60 s was used to collect 512 × 512 pixels (200 μm × 200 μm) fluorescence lifetime images simultaneously for NAD(P)H and the fluorescently labeled antibodies, followed by collection of FAD. A daily instrument response function (IRF) was collected from the second harmonic generation of a urea crystal. Images were collected for 3-10 fields of view for each cell type, depending on cell type abundance.

### 1.3 Image Analysis

Pixel-level photon arrival time histograms were extracted using SPCImage (v8.9, Becker & Hickl). To enhance the accuracy of fluorescence lifetime measurements, images were binned using a bin factor of two (summing 25 pixels in a 5×5 pixel kernel) for NAD(P)H images and a bin factor of three (summing 49 pixels in a 7×7 pixel kernel) for FAD images to reach a minimum of 2,500 photons in each decay. Low-intensity background pixels were removed through thresholding. NAD(P)H and FAD exist in free and protein-bound conformations with distinct short and long lifetime components [49,50]. Using an iterative IRF reconvolution and parameter optimization to maximize the likelihood of the observed data under Poisson statistics/distribution (Maximum Likelihood Estimation algorithm), the pixel-wise NAD(P)H and FAD decay curves were fit to a biexponential model (**Eq 1**).

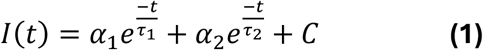

Where I(t) is the intensity of autofluorescence at time t after the laser pulse. The short (quenched) and long (unquenched) lifetime components are given as τ_1_ and τ_2_, respectively, while the amplitude of each component is given as α_1_ and α_2_, respectively. C accounts for non-decaying background light amplitude. Additionally, the mean lifetime was calculated at each pixel for NAD(P)H and FAD (**Eq 2**).

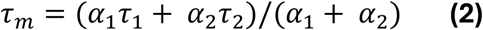

Whole-cell masks were generated from the antibody fluorescence intensity images using the cyto2 model in Cellpose [67] with diameter = 50 pixels (∼20 µm), cell probability threshold = −0.50, and flow threshold = −25. Masks were overlayed on the NAD(P)H intensity images and manually corrected using Napari [68] to remove masked debris and motion artifacts, ensuring masks cover only single cells. Fluorescence lifetime variables were averaged across each single-cell mask in Python using cell-analysis-tools [69]. In the results, α_1_ and α_2_ refer to the normalized fractions α_1_ / (α_1_ + α_2_), and α_2_ / (α_1_ + α_2_), respectively, and are given as percentage values. The optical redox ratio (ORR) was calculated at the cell level using the summed NAD(P)H and FAD fluorescence over all cell pixels using **Equation 3**, which constrains the ORR between 0 and 1.

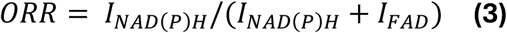

Cells with low photon counts (summed NAD(P)H average cell intensity < 10 a.u.), small masks unlikely to be cells (<70 pixels), and cells with poor goodness of fit (Х^2^>1.3) or NAD(P)H τ_1_<350 ps were not included in the analysis. Single variable plots were generated using matplotlib [70] and seaborn [71].

### 1.4 24-hour Activation and Metabolite Analysis

After 24 hours in culture, media from the unstained activated or quiescent PBMC wells was collected and frozen at −20°C. Cells were resuspended in 200 μL fresh media with 0.10 μL 10 mg/mL Hoechst 33342 (ThermoFisher) and 1 μL PerCP-αCD69 antibody (**Supp Table 1**). After 30 minutes, excess antibody was washed as described above and cells were resuspended in 200 μL media. Intensity images were collected on a Nikon Ti-2E coupled to a 380 to 660 nm SOLA light engine (Lumencor). Light was collected through a 40X (NA = 0.95) air objective (Plan Apo, Nikon) and detected by a Hamamatsu Orca-Flash digital CMOS camera. Hoechst was excited for 1 ms from 340-380 nm at 26% power (0.56 W/cm^2^) and emission was detected from 435-485 nm. PerCP-αCD69 was excited for 20 ms from 550-590 nm at 26% power (3.3 W/cm^2^) and emission was detected from 608-683 nm.

Single cells were identified from brightfield images using the cyto2 model in Cellpose with diameter = 100, cell probability threshold = −0.50, and flow threshold = −25. Masks were applied to the PerCP-αCD69 images and an average PerCP intensity was collected for each cell, from which a histogram was constructed in seaborn/matplotlib showing CD69- and CD69+ populations. The CD69+ threshold was set at 1.5 standard deviations above the mean of the quiescent population.

A lactate assay was performed to verify upregulation of aerobic glycolysis with activation [72]. The lactate colorimetric assay was carried out according to the assay kit (BioVision). Briefly, 2 μL of each media sample was added to 198 μL assay buffer and 50 μL of solution was added to three replicate wells of a 96 well plate. 50 μL of reaction mix (2 μL probe, 2 μL enzyme mix, and 46 μL assay buffer) was added to each well and left to incubate for 30 minutes at 37°C. The optical density (OD) was measured at 570 nm using a microplate reader. A standard curve was fit to the results of a concentration series of the lactate standard using an ordinary least squares regression model with an R-squared of 0.9975.

### 1.5 Classifiers and Statistics

Random forest classifiers were trained in python using scikit-learn [73] on a random selection of 70% of the cells and tested on the remaining 30% using 10 OMI variables (NAD(P)H and FAD τ_m_, τ_1_, τ_2_, α_1_, the ORR, and cell area). Strength of the classifiers was assessed using the receiver operating characteristic (ROC) curve, area under the curve (AUC) of the ROC, accuracy, precision, and recall. Classifiers were trained and tested on different random sets of cells to check for consistency in these metrics. To prevent model bias toward majority classes and ensure that misclassifications in any category were penalized equally, class balancing was performed prior to training by randomly undersampling all categories to match the frequency of the minority class.

Single cell heterogeneity was visualized with Uniform Manifold Approximation and Projection (UMAP) [74] plots generated in Python using Scikit-learn using 10 OMI variables (NAD(P)H and FAD τ_m_, τ_1_, τ_2_, α_1_, the ORR, and cell area). UMAPs were computed using euclidean distance, the nearest neighbor parameter was set to 6, and the minimum distance was set to 0.6. Heterogeneity analysis was performed by computing mean and standard deviation for each variable of each activated or quiescent cell type. The coefficient of variation was calculated as the ratio of the standard deviation to the mean, providing a normalized measure of single-cell heterogeneity across different variables. The Z-score heatmap was constructed in python using the seaborn clustermap function. Single cells were hierarchically clustered based on OMI variables using Ward’s linkage with Euclidean distance. Activation condition, donor, and cell type were added as row annotations and were not included in the clustering analysis.

Unless otherwise stated, statistical analysis of single variables was performed using Cohen’s d [75], a measure of effect size, due to the large number of single cell data points [76] (**Eq 4**).

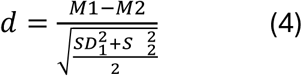

Where M and SD represent the mean and standard deviation of each group, respectively. Values where |d|<0.2 represent no change, 0.2≤|d|<0.5 represent a small change, 0.5≤|d|<0.8 represent a medium change, and |d|≥0.8 represent a large change [76]. To reduce sampling error, a minimum of 100 cells were assessed per cell type in each condition.

## Results

### 1.1 OMI classifies quiescent PBMC subtypes

Quiescent PBMCs were stained using fluorescent-labeled antibodies, enabling identification of the following cell types within the heterogeneous PBMC culture: helper T cells (CD4+), cytotoxic T cells (CD8+), monocytes (CD14+), B cells (CD19+), NK cells (CD56+). Notably, monocytes displayed a low NAD(P)H τ_m_, high NAD(P)H α_1_, and large cell area in the quiescent state compared to other cell types [**Fig. 2(a)-2(d)**]. Resting B cells also displayed a slightly higher NAD(P)H α_1_ when compared to T cells and NK cells [**Fig. 2(c)**]. All other OMI variables for quiescent PBMCs are plotted by subtype in **Supp. Fig. 1**.

**Figure 2.**
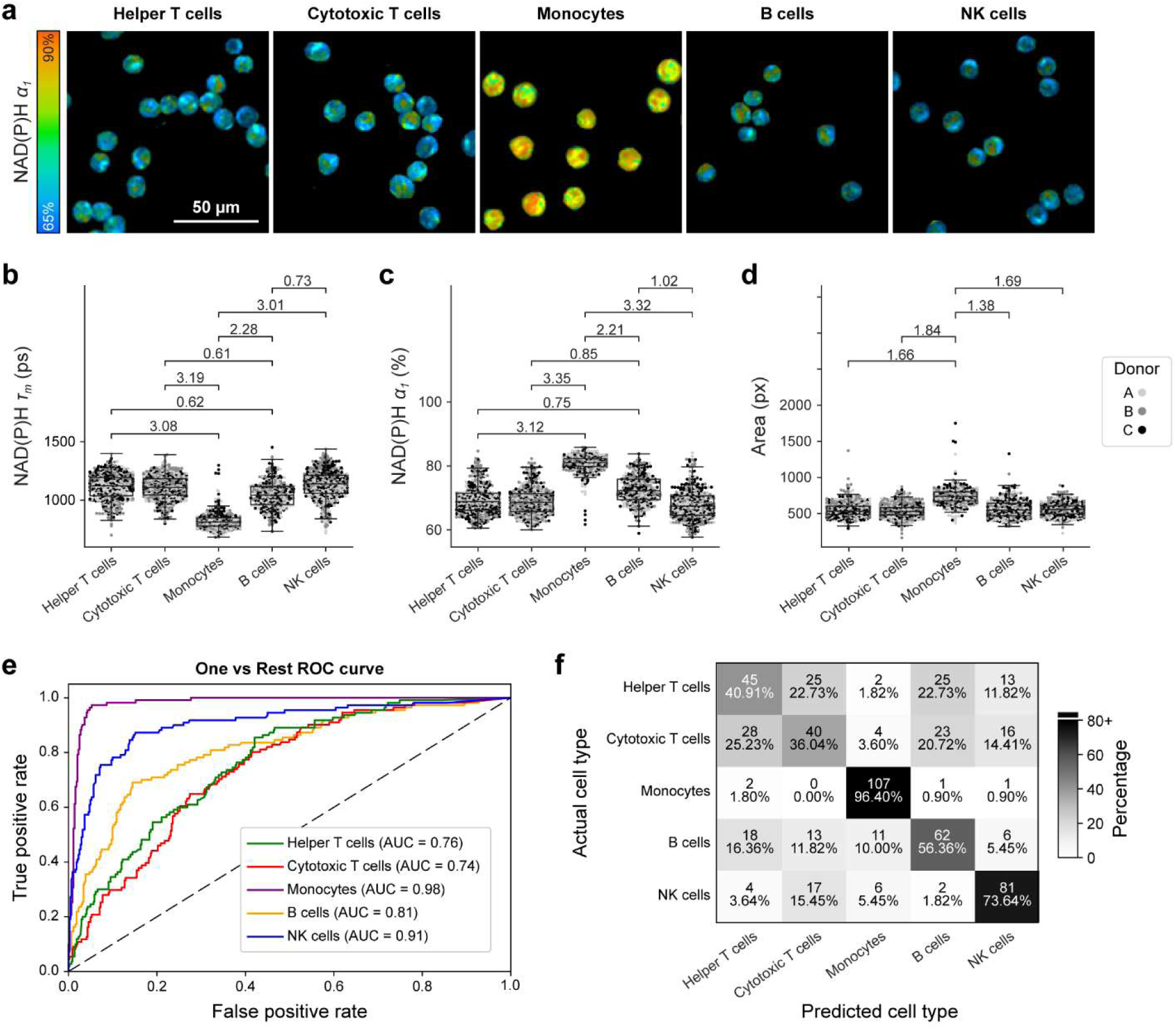
Quiescent monocytes and lymphocytes have distinct autofluorescence. (a) Representative images of quiescent PBMC cell populations. (b-d) NAD(P)H τ_m_, NAD(P)H α_1_, and cell area for quiescent PBMC cell populations. Data are displayed as box-and-whisker plots, representing the median and interquartile range (IQR), with whiskers at 1.5*IQR. Plots are overlaid on individually plotted cells shaded by donor, A, B, or C. Effect size given as absolute value of Cohens d. Insignificant and small effect sizes (d<0.5) not shown, medium (0.5≤|d|<0.8), large (|d|≥0.8). (e-f) A one versus rest random forest classifier was trained on 70% of data and tested on the remaining 30% using all OMI variables (NAD(P)H and FAD τ_m_, τ_1_, τ_2_, and α_1_, the ORR, and cell area). (e) ROC and (f) confusion matrix for classification of each cell type showing cell number and percent of cells across a row. n = 553 helper T cells, 554 cytotoxic T cells, 425 monocytes, 368 B cells, 510 NK cells.

A one-versus-rest random forest classifier using 10 OMI variables was trained on 70% of the quiescent PBMCs and tested on the remaining 30%. Monocytes were classified effectively (AUC = 0.98, Recall = 96.4%) [**Fig. 2(e)-2(f)**]. The variables that most effectively discriminated monocytes from the lymphocytes were NAD(P)H τ_m_ (feature importance = 57.3%) and NAD(P)H α_1_ (feature importance = 18.0%) [**Supp. Fig. 2**]. Cytotoxic T cells, helper T cells, and B cells were not strongly resolved based on OMI variables [**Fig. 2(e)-2(f)**]. However, classification of NK cells was moderately accurate (AUC=0.91, Recall = 68.9%) [**Fig. 2(e)-2(f)**] with the most important variables for classification being ORR (feature importance = 20.5%), FAD τ_1_ (feature importance = 18.6%), and FAD α_1_ (feature importance = 16.3%) [**Supp. Fig. 2**].

### 1.2 OMI classifies early activation in PBMC subtypes

To capture changes with activation, PBMCs were plated and cultured with potent immune cell activators (40 nM PMA and 1 μM ionomycin) for 2 hours before imaging. Representative images demonstrate qualitative changes with early activation including the greater intracellular heterogeneity and granularity observed in the monocytes [**Fig. 3(a)**]. As in the quiescent condition, activated monocytes had lower NAD(P)H τ_m_, higher NAD(P)H α_1_, and larger cell areas than other activated PBMCs [**Fig. 3(b)-3(d)**]. However, effect sizes between monocytes and other PBMC cell types were smaller when activated [**Fig. 3(b)-3(d)**] compared to quiescence [**Fig. 2(b)-2(d)**], demonstrating increased metabolic homogeneity with activation. All other OMI variables for quiescent PBMCs are plotted by subtype in **Supp. Fig. 3**.

**Figure 3.**
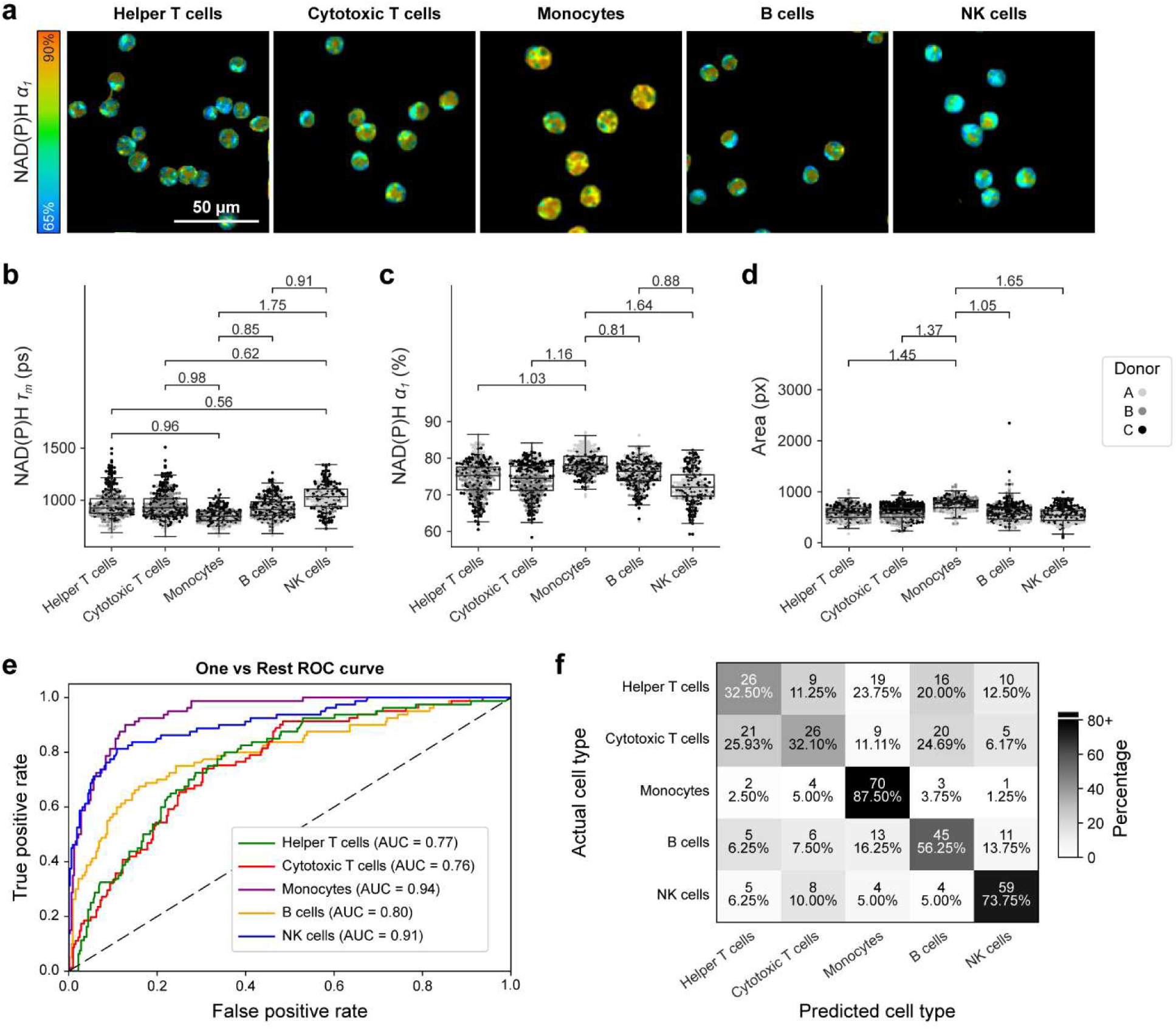
Activated monocytes and lymphocytes have distinct autofluorescence. (a) Representative images of activated (2-hour PMA/Ionomycin) cell populations within PBMCs. (b-d) NAD(P)H τ_m_, NAD(P)H α_1_, and cell area for activated PBMC cell populations. Data are displayed as box-and-whisker plots, representing the median and interquartile range (IQR), with whiskers at 1.5*IQR. Plots are overlaid on individually plotted cells shaded by donor, A, B, or C. Effect size given as absolute value of Cohens d. Insignificant and small effect sizes (d<0.5) not shown, medium (0.5≤|d|<0.8), large (|d|≥0.8). (e-f) A one versus rest random forest classifier was trained on 70% of data and tested on the remaining 30% using all OMI variables (NAD(P)H and FAD τ_m_, τ_1_, τ_2_, and α_1_, the ORR, and cell area). (e) ROC and (f) confusion matrix for classification of each cell type showing cell number and percent of cells across a row. n = 363 helper T cells, 353 cytotoxic T cells, 305 monocytes, 340 B cells, 267 NK cells.

A one-versus-rest random forest classifier using 10 OMI variables was trained on 70% of the activated PBMCs and tested on the remaining 30%. Monocytes remained distinct from the lymphocytes within the activated PBMCs (AUC = 0.94, Recall = 87.5%) [**Fig. 3(e)-3(f)**], though had slightly weaker predictive power than in the quiescent PBMCs (AUC = 0.98, Recall = 96.4%) [**Fig. 2(e)-2(f)**]. In the activated PBMCs, monocyte classification depended primarily on cell area (feature importance = 31.0%), followed by NAD(P)H α_1_ (feature importance = 15.6%) [**Supp. Fig. 4**]. Cytotoxic T cells, helper T cells, and B cells were not clearly identifiable at this early activation time point. Activated NK cells retained moderate classification accuracy (AUC = 0.91, Recall = 73.8%) [**Fig. 3(e)-3(f)**] with highest variable weights from FAD τ_1_ (feature importance = 21.1%) and NAD(P)H τ_m_ (feature importance = 16.1%) [**Supp. Fig. 4**].

**Figure 4.**
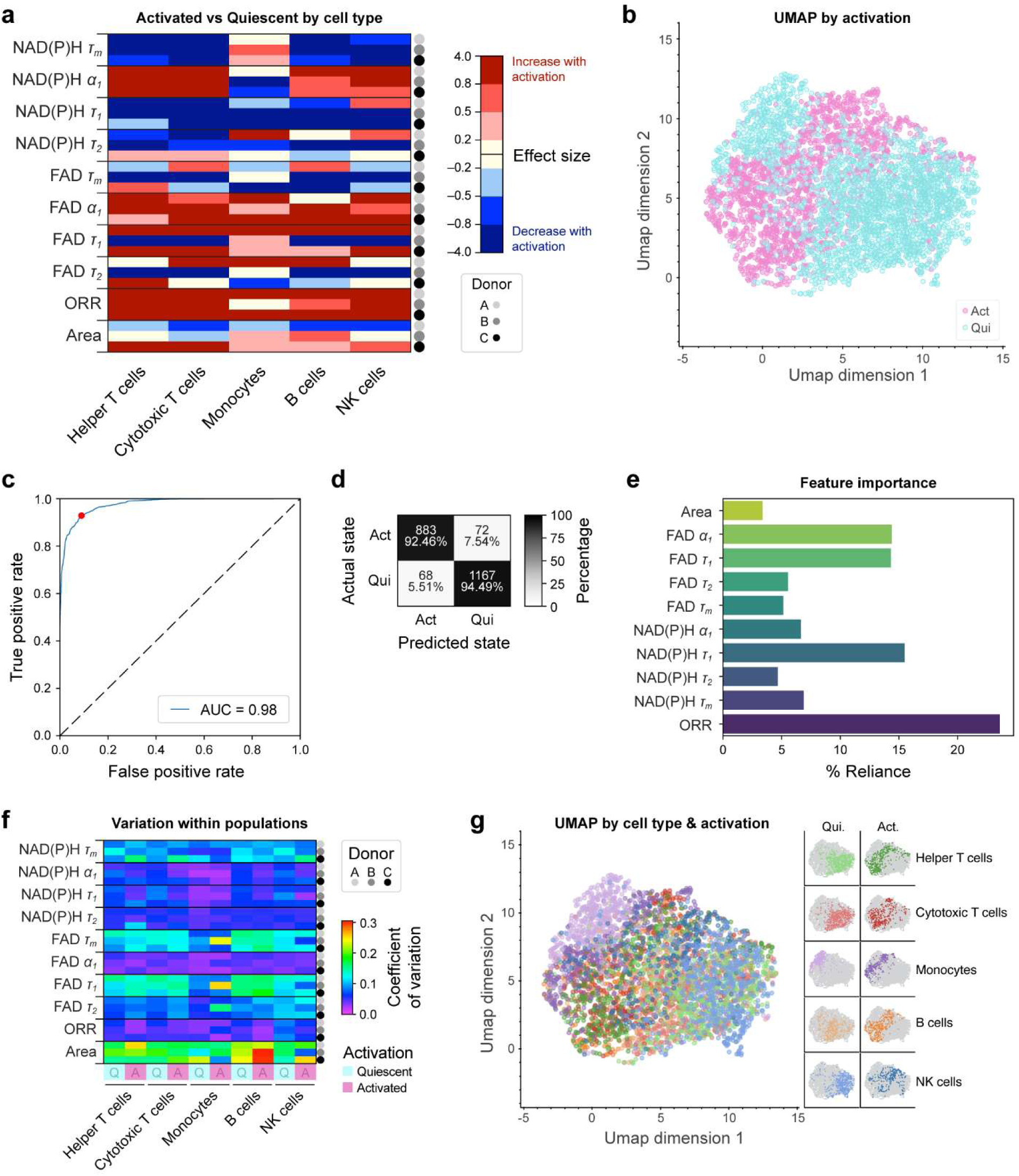
Metabolic Heterogeneity increases with activation. (a) Effect size of activated vs quiescent cell types by donor and variable colored by low (0.2≤|d|<0.5), medium (0.5≤|d|<0.8), large (|d|≥0.8), and nonsignificant (|d|<0.2) effect sizes, and directionality of change with activation: Increasing (red) and decreasing (blue). (b) UMAP displays clustering by activation condition. (c-d) Random forest classifier ROC curve with plotted operating point and confusion matrix of activated vs quiescent PBMCs showing cell number and percent of cells across a row (Accuracy = 92.7%). (e) Relative importance of the 10 OMI variables in the random forest classifier. (f) Coefficient of variation (COV = standard deviation / mean) displays metabolic variability within each quiescent and activated cell population. (g) UMAP displays clustering by activation condition and cell type. n = 2410 quiescent PBMCs and 1628 activated PBMCs. Cell counts by donor, cell type, and activation can be found in supplemental table 2.

Activation was confirmed later, after 24 hours in culture, demonstrating increased lactate production (70% increase) and CD69 expression (79.6% CD69+) compared to quiescent PBMCs (<1% CD69+) [**Supp. Fig. 5**], consistent with existing literature [63, 64].

### 1.3 OMI assesses single-cell metabolic heterogeneity in PBMCs

Cell type dependent activation dynamics are summarized in **Figure 4(a)**, which shows the effect size between the quiescent and activated condition for each variable and cell type (cell counts available in **Supp. Table. 2**). All lymphocytes (T cells, B cells, NK cells) had decreased NAD(P)H τ_m_ and increased NAD(P)H α_1_ at 2 hours post-activation compared to the pre-activation (quiescent) condition, which is consistent with prior findings with immune activation [13, 63, 64] [**Fig. 4(a)**]. Conversely, monocytes showed increased NAD(P)H τ_m_ and decreased NAD(P)H α_1_ at 2 hours post-activation compared to the quiescent condition [**Fig. 4(a)**]. All PBMC subtypes generally increased ORR with activation, demonstrating a shift towards a reduced intracellular environment, and had decreased NAD(P)H τ_1_ [**Fig. 4(a)**].

Separate clusters of activated and quiescent PBMCs can be observed using uniform manifold approximation projection (UMAP) of 10 OMI variables [**Fig. 4(b)**]. Bulk quiescent and activated PBMCs can be classified with high accuracy (AUC = 0.98, Accuracy = 93.6%) [**Fig. 4(c)-4(d)**]. The variables with the highest weights in this classifier for activation in PBMCs include the ORR (feature importance = 23.6%) and NAD(P)H τ_1_ (feature importance = 15.5%) [**Fig. 4(e)**]. These variables also displayed the most consistent shifts with activation across all PBMC subtypes [**Fig. 4(a)**].

Variation of OMI variables within each cell type was assessed using the coefficient of variation (COV) [**Fig. 4(f)**]. While variables like NAD(P)H α_1_, FAD α_1_, and ORR had low levels of variation in both quiescent and activated conditions, most cell types and variables increased in heterogeneity with activation. FAD τ_1_, FAD τ_m_, and cell area all demonstrated the greatest heterogeneity with activation, though these variables already displayed relatively high levels of heterogeneity in quiescence [**Fig. 4(f)**].

When colored by cell and activation, the UMAP displays distinct clusters of quiescent and activated monocytes (purple) and NK cells (blue) [**Fig. 4(g)**]. Helper T cells (green), cytotoxic T cells (red), and B cells (orange) qualitatively cluster together though the activated cluster of these cells is distinct from the quiescent cluster [**Fig. 4(g)**]. Notably, monocytes occupy a distinct cluster that is more closely associated with the activated PBMCs, which is further recapitulated using hierarchical clustering analysis of the same 10 OMI variables [**Supp. Fig. 6**]. Due to size constraints, the heatmap was broken along the first two hierarchical clusters. Cluster 1 is defined by low NAD(P)H α_1_ and primarily contains quiescent lymphocytes while cluster 2 displays high NAD(P)H α_1_ and contains activated PBMCs and quiescent monocytes.

NK cells are canonically identified using CD56, which is expressed at high (bright) and low (dim) levels, where CD56 bright NK cells account for approximately 10% of the blood NK cell population, produce more cytokines, are less cytotoxic, and have greater reliance on glycolysis than the CD56 dim population [22,77]. Therefore, we explored whether OMI could identify metabolic differences in CD56 dim vs. bright NK cells. We defined CD56 bright NK cells with a threshold defined at 100 a.u., which captured approximately the top 10% brightest αCD56 NK cells [**Supp. Fig. 7(a)-7(b)**]. Significant decreases in NAD(P)H τ_m_, τ_1_, and τ_2_ were observed in the CD56 bright vs. dim cells [**Supp. Fig. 7(c)-7(f)**], consistent with increased glycolysis in the CD56 bright population, with some clustering on a UMAP of all OMI variables [**Supp. Fig. 7(g)**].

## Discussion

PBMCs are routinely assessed for disease diagnostics and cell therapy development but traditional tools do not consider metabolic heterogeneity, which provides critical functional information. For example, OMI previously identified metabolic subsets of CAR T cells that were not apparent with surface marker expression alone, which lead to improved tumor cures *in vivo* [13]. Similarly, metabolic flux analysis of patient PBMCs identified early markers sepsis, which can guide timely treatment decisions to improve patient survival [37]. Here, we demonstrate a label-free and non-destructive imaging technique to analyze single-cell metabolic phenotypes within quiescent and activated primary human PBMCs. Our analysis revealed that OMI can identify monocytes within the quiescent and activated PBMCs (AUC > 0.94) and NK cells within quiescent and activated PBMCs (AUC = 0.91) with relatively high classification accuracy. Additionally, OMI can accurately classify early activation state within PBMCs (2 hours post-stimulation, AUC = 0.98). While CD69 and CD25 are well established markers of immune activation, here we demonstrate a label-free approach that could enable rapid and automated analysis in a clinical workflow because personnel time is not needed for the labeling steps.

Classification accuracy for most cell subtypes diminished within the activated PBMCs compared to the quiescent PBMCs [**Figs. 2(e-f) and 3(e-f)**]. This could be due to similar early metabolic changes across PBMCs with PMA/ionomycin-induced activation, which indiscriminately upregulates metabolic demand in all PBMCs via protein kinase C activation. However, more physiologically relevant immune activators (e.g., antibodies, interleukins, and other secreted factors) would likely produce different metabolic and timed responses within PBMC subsets and may require training new classifiers.

Our findings with PBMC activation are consistent with prior studies that found an increase in NAD(P)H α_1_ and the ORR with activation of T, B, and NK cells in monoculture at various time points post-activation using varying activation methods [13, 62–64]. While activation is traditionally confirmed using surface marker expression, particularly using anti-CD69 or anti-CD25, these markers are imperfect and can take several hours to fully express, while secreted factors and metabolites can require several hours to accumulate to detectable levels within media followed by additional processing times. This highlights the speed and sensitivity of OMI to early activation (2 hours post-stimulation). Additionally, previous reports demonstrate that innate myeloid cells (e.g., monocytes) have higher glycolytic demand than lymphocytes [18], which supports clustering of monocytes with activated lymphocytes in our UMAP [**Fig. 4(g)**] and hierarchical clustering [**Supp. Fig. 6**].

The universal decrease in NAD(P)H τ_1_ with activation suggests alterations in the biochemical environment of each cell subtype that promotes quenching of free NAD(P)H, consequently shortening the fluorescence lifetime. Ionomycin is a potent calcium ionophore and thus increases intracellular calcium concentrations that may promote NAD(P)H quenching. While our observed NAD(P)H τ_1_ decrease is large, prior studies have reported decreases in NAD(P)H τ_1_ with more physiologically relevant immune activators including tetrameric antibody against CD2/CD3/CD28 in T cells and IL-12/IL-15/IL-18 stimulation of NK cells, demonstrating that this effect is not uniquely induced by ionomycin [13, 63, 64].

Here, no significant differences were observed between CD4+ and CD8+ T cells, which have previously shown separation in OMI parameters within CD3+ T cell systems [63]. This discrepancy is likely due to the multiple cell subtypes within the PBMC samples, where secreted factors and cell-cell interactions influence cell metabolism compared to monocultures of identical cells. Additionally, within the T cell cultures previously described, composition of the T cell mixtures was demonstrated to affect metabolic features [63]. Furthermore, these prior studies used a tetrameric antibody against CD2/CD3/CD28, which induces a more physiologically relevant activation response in these cells. This response is known to differ between CD4+ and CD8+ T cells while PMA/ionomycin produces more similar responses in both CD4+ and CD8+ T cells [63, 78]. This highlights the critical roles that both the activator and the microenvironment play in modulating immune cell metabolism.

Cell metabolism provides a rapid and sensitive measurement of immune cell function that is difficult to capture with surface marker expression alone. Therefore, single-cell measurements of metabolism within PBMCs could enhance current diagnostic tests (e.g., sepsis and SLE) and quality screening for cell therapy (e.g., CAR T cell and tumor infiltrating lymphocyte therapy). While traditional measurements of metabolism require labels or destroy the sample, OMI provides a label-free and non-destructive measurement of single-cell metabolic features that can discriminate metabolic subsets of PBMCs. Therefore, OMI can be multiplexed with traditional destructive methods like single-cell RNA sequencing and antibody-labeled flow cytometry, or OMI could be used independently to characterize cells before their use in downstream applications of cell therapies. Together, this study demonstrates that OMI successfully characterizes cell heterogeneity within complex cultures, identifying monocytes and NK cells within PBMCs and accurately classifying early activation status within PBMCs. NAD(P)H lifetime measurements can further be performed in a flow geometry [79], which will streamline translation to clinical and biomanufacturing settings for immunophenotyping in disease monitoring, drug screens, and the development of cell therapies.

## Conclusion

PBMCs are clinically used to monitor disease state and as a starting material for cell therapies. Cell metabolism is a sensitive feature of PBMC function and quality, but there are currently no non-destructive techniques to measure single-cell metabolism within PBMCs. Here, we find that OMI provides a label-free measurement of single-cell metabolism that identifies immune cell subsets (monocytes, NK cells) and early activation state (2 hours post-stimulation) within PBMCs without the need for staining, enabling high throughput metabolic-screens. OMI could therefore provide additional metabolic information to complement traditional measurements of PBMCs, which could improve disease monitoring, the development of immune therapies, and other applications where touch-free metabolic phenotyping is beneficial.

## Code and Data Availability

The authors declare that all relevant data supporting the findings of this study are available within the article and its Supplementary Material. All Excel files used to generate figures and support conclusions and the associated codes have been deposited in the following GitHub repository: https://github.com/skalalab/Riendeau_PBMC_OMI. Raw images are available from the corresponding authors upon reasonable request.

## Acknowledgments

We thank Matthew Stefely for assisting in generating figures. This work was supported by the National Institutes of Health (R01 CA278051, R01 HL165726) and the NSF Engineering Research Center for Cell Manufacturing Technologies (CMaT) (Grant No. 1648035).

