## Supplemental Material for "Autofluorescence lifetime imaging resolves cell heterogeneity within peripheral blood mononuclear cells"

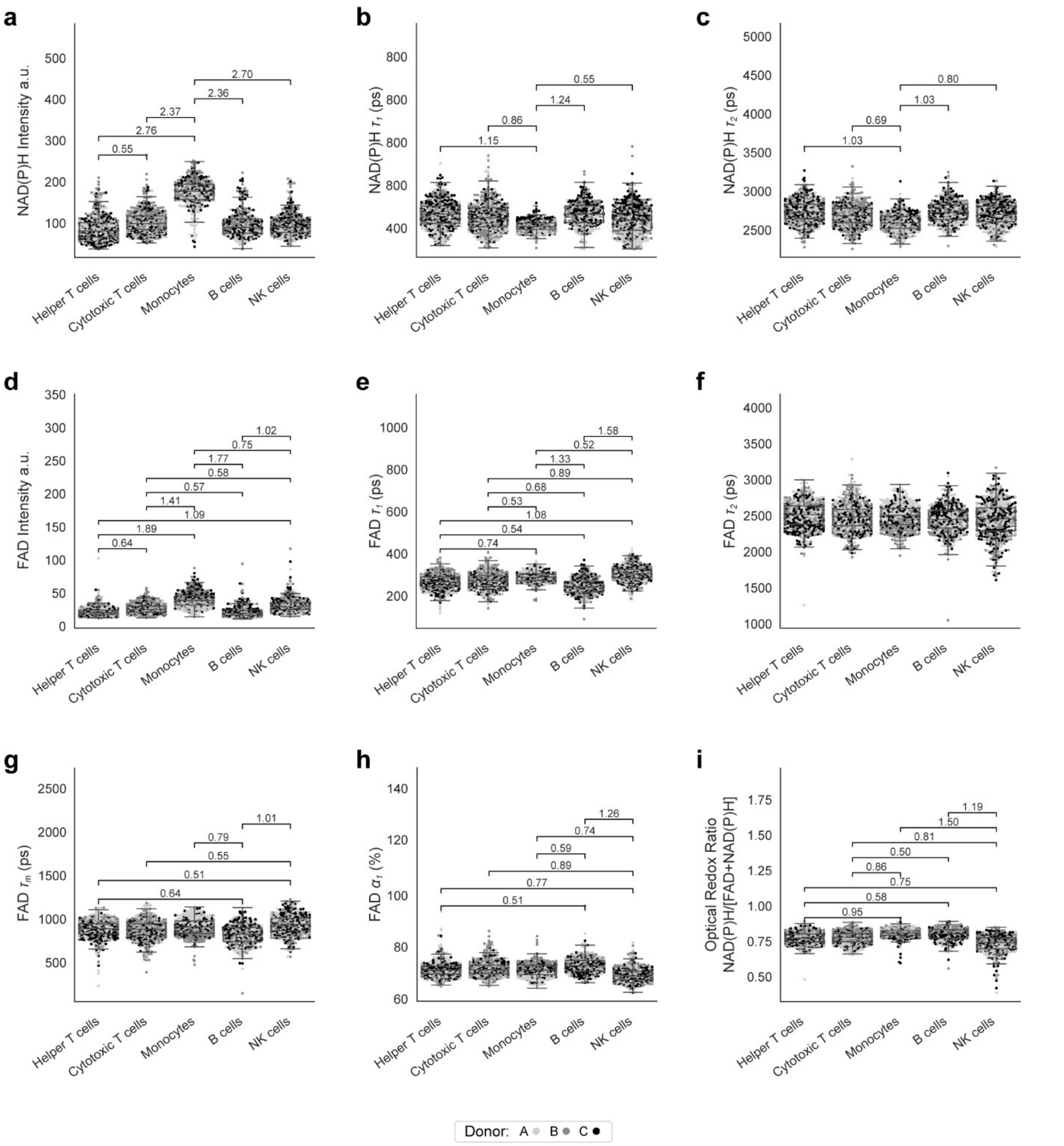

**Supplemental Figure 1. OMI features of quiescent PBMC immune populations.** (a-c) NAD(P)H features plotted by cell type. (d-h) FAD features plotted by cell type. (i) Optical redox ratio plotted by cell type. Data are displayed as box-and-whisker plots, representing the median and interquartile range (IQR), with whiskers at 1.5*IQR. Plots are overlaid on individually plotted cells colored by donor, A, B, or C. n = 553 helper T cells, 554 cytotoxic T cells, 425 monocytes, 368 B cells, 510 NK cells. Effect size given as absolute value of Cohens d. Insignificant and small effect sizes (|d|<0.5) not shown, medium (0.5≤|d|<0.8), large (|d|≥0.8).

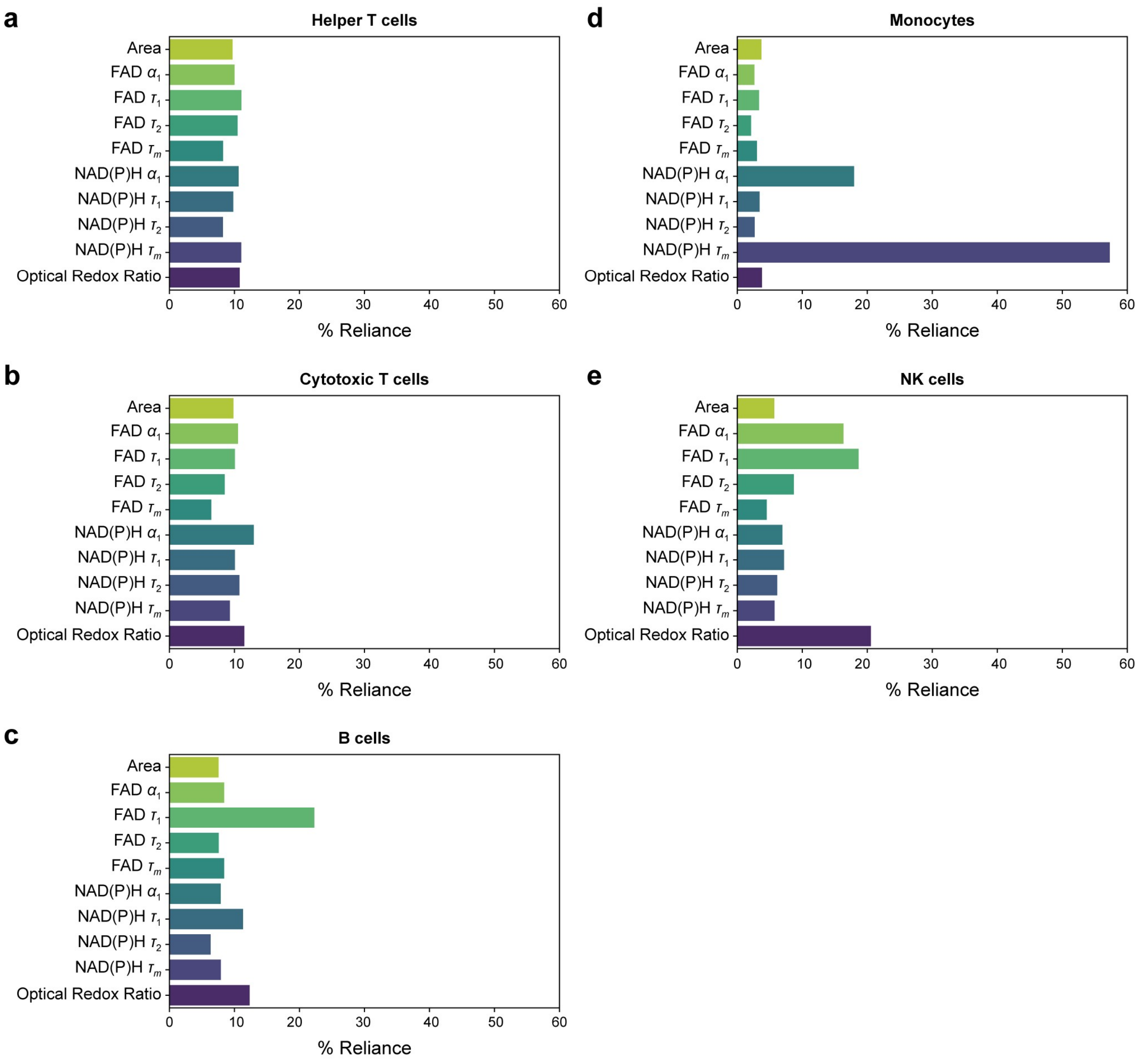


**Supplemental Figure 2. Quiescent PBMC one versus rest random forest classification feature weights.** Relative importance of the 10 OMI variables for classification of (a) helper T cells, (b) cytotoxic T cells, (c) B cells, (d) monocytes, and (e) NK cells from the rest of the PBMC immune populations.

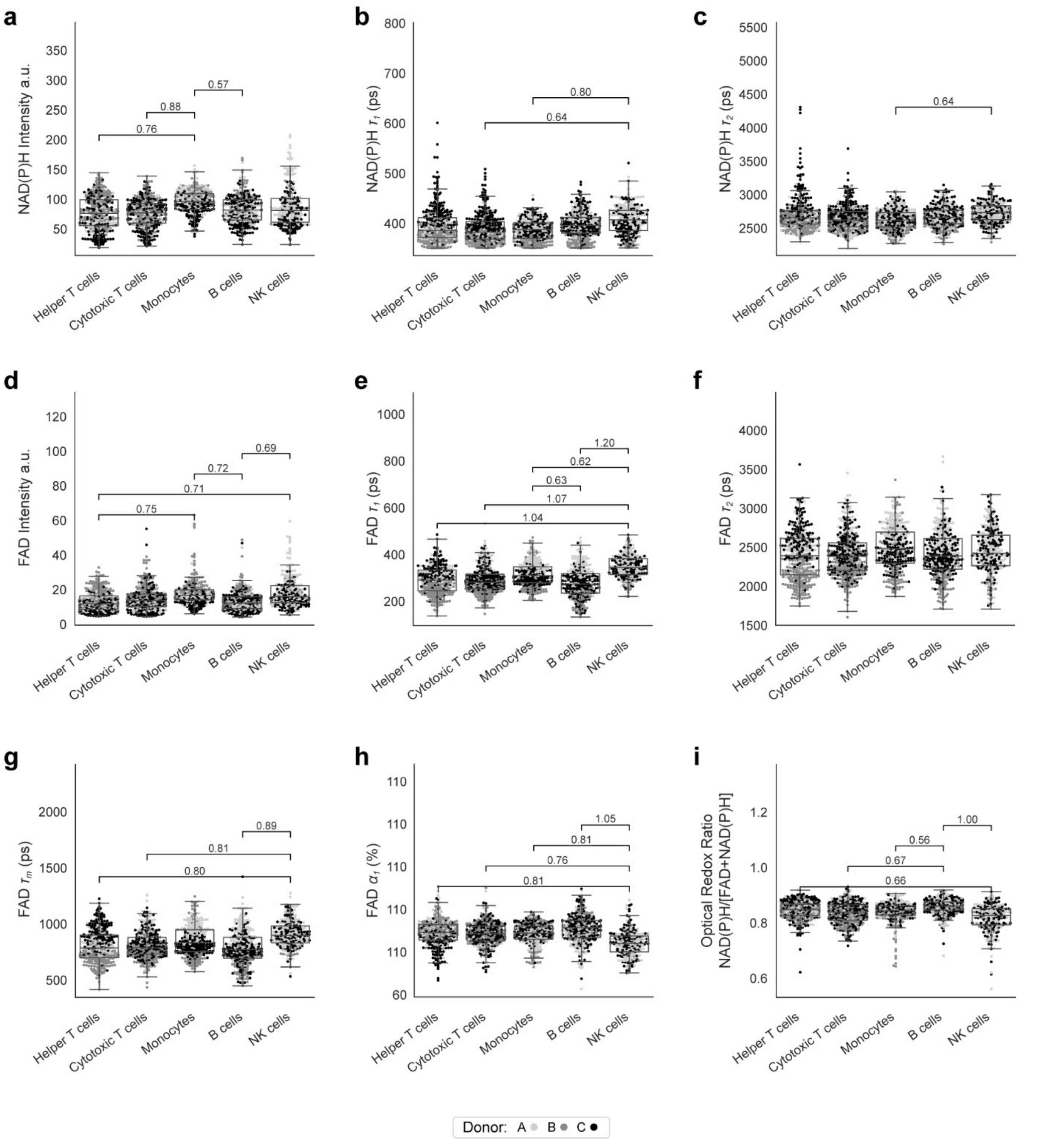


**Supplemental Figure 3. OMI features of PMA and Ionomycin-activated PBMC immune populations.** (a-c) NAD(P)H features plotted by cell type. (d-h) FAD features plotted by cell type. (i) Optical redox ratio plotted by cell type. Data are displayed as box-and-whisker plots, representing the median and interquartile range (IQR), with whiskers at 1.5*IQR. Plots are overlaid on individually plotted cells colored by donor, A, B, or C. n = 363 helper T cells, 353 cytotoxic T cells, 305 monocytes, 340 B cells, 267 NK cells. Effect size given as absolute value of Cohens d. Insignificant and small effect sizes (|d|<0.5) not shown, medium (0.5≤|d|<0.8), large (|d|≥0.8).

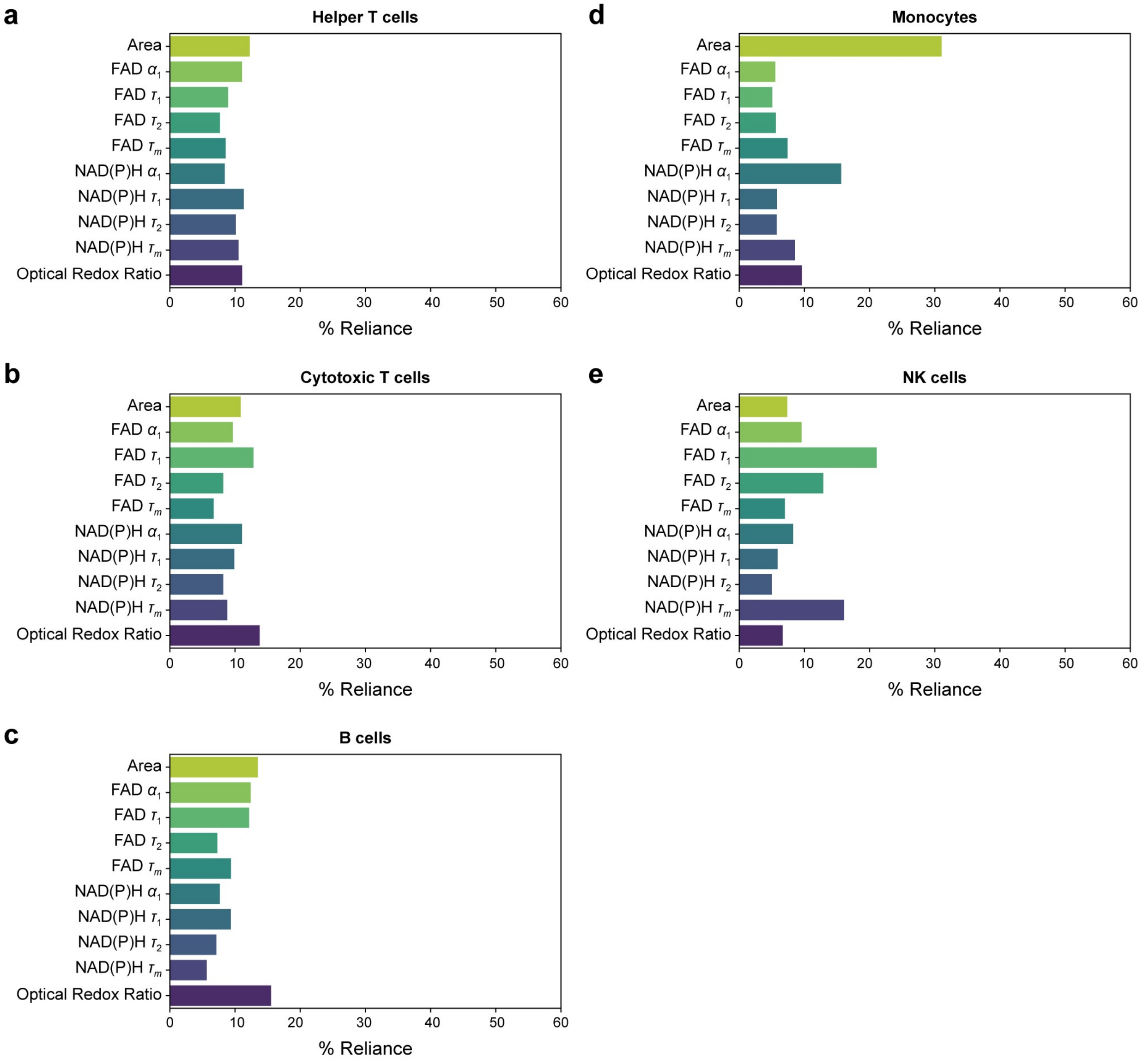


**Supplemental Figure 4. PMA and Ionomycin-activated PBMC one versus rest random forest classification feature weights.** Relative importance of the 10 OMI variables for classification of (a) helper T cells, (b) cytotoxic T cells, (c) B cells, (d) monocytes, and (e) NK cells from the rest of the PBMC immune populations.

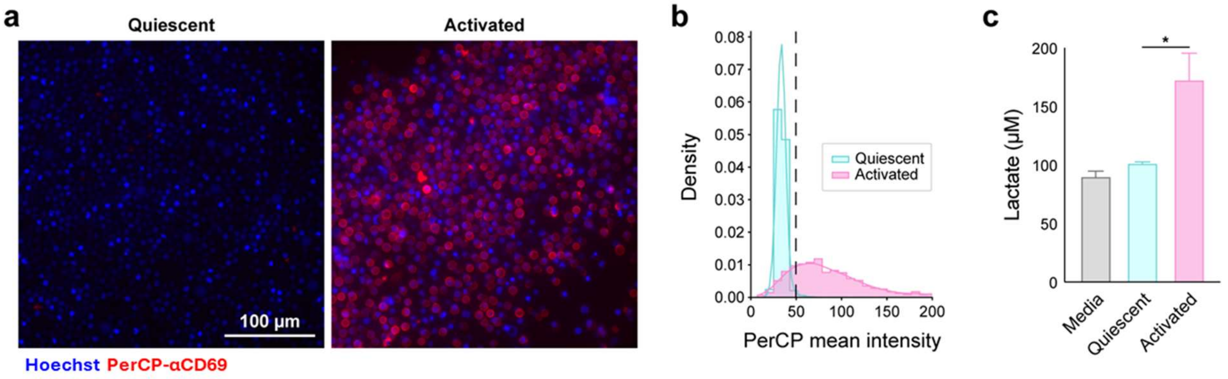


**Supplemental Figure 5. 24-hour quiescent and PMA/Ionomycin-activated PBMCs.** (a) Representative Hoechst (blue) and αCD69 (red) staining. (b) Quantification of single-cell staining reveals 79.6% PerCP-αCD69 expression after 24 hours in activating conditions. Threshold defined at 1.5 standard deviations over the mean PerCP-αCD69 intensity of the quiescent group. (c) Lactate production after 24 hours was increased by 70% with activation compared to quiescent. n= 3153 activated and 1839 quiescent PBMCs. *P<0.05, two-tailed unpaired T-test.

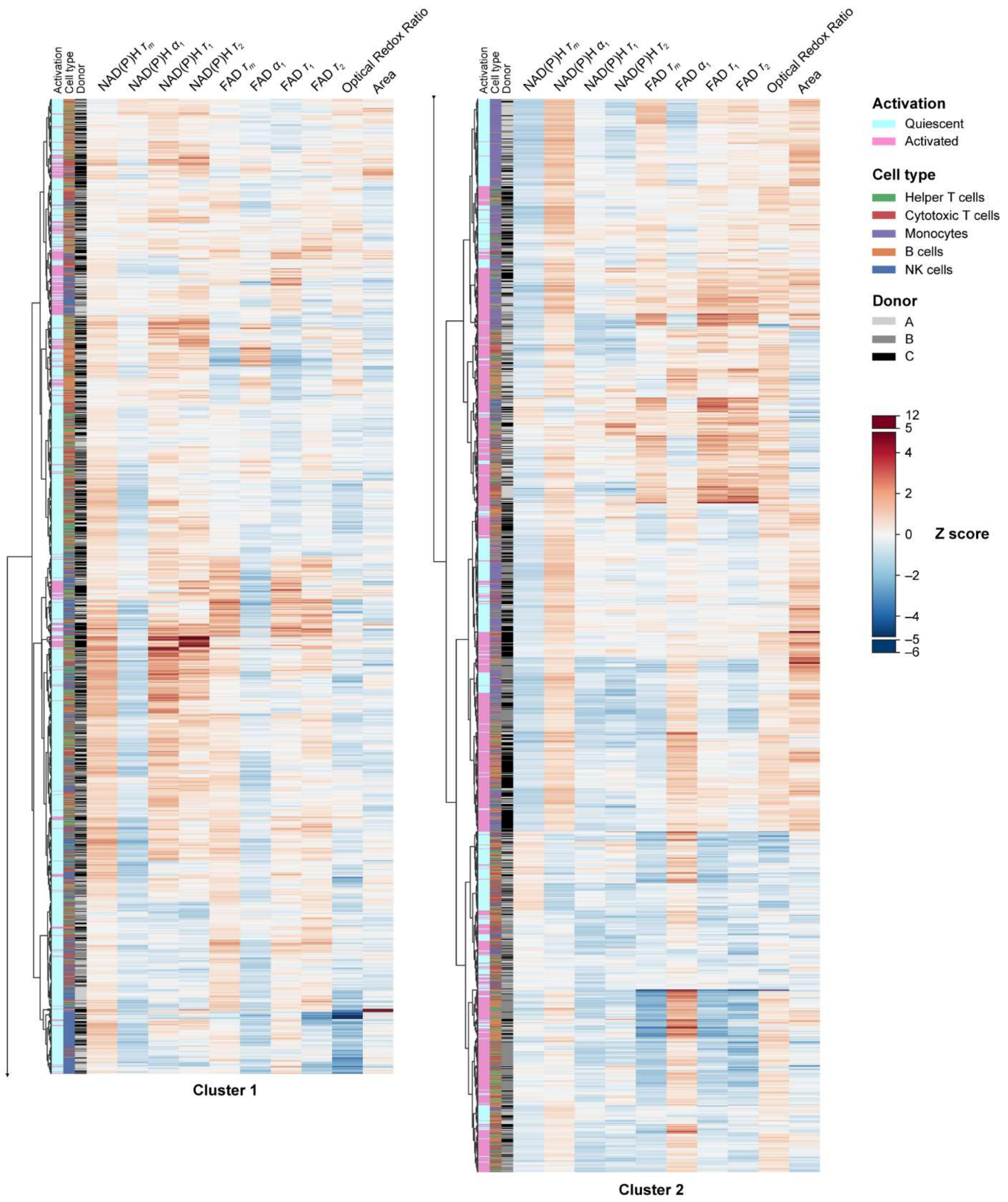


**Supplemental Figure 6. Hierarchical clustering of all PBMCs**. Heatmap and hierarchical clustering of single-cells was calculated based on the z-scores (the difference between cell mean and population mean divided by the population standard deviation) of 10 OMI variables (NAD(P)H and FAD $\tau_m$, $\tau_1$, $\tau_2$, $\alpha_1$, the ORR, and cell area). Cell counts by donor, cell type, and activation can be found in supplemental table 2.

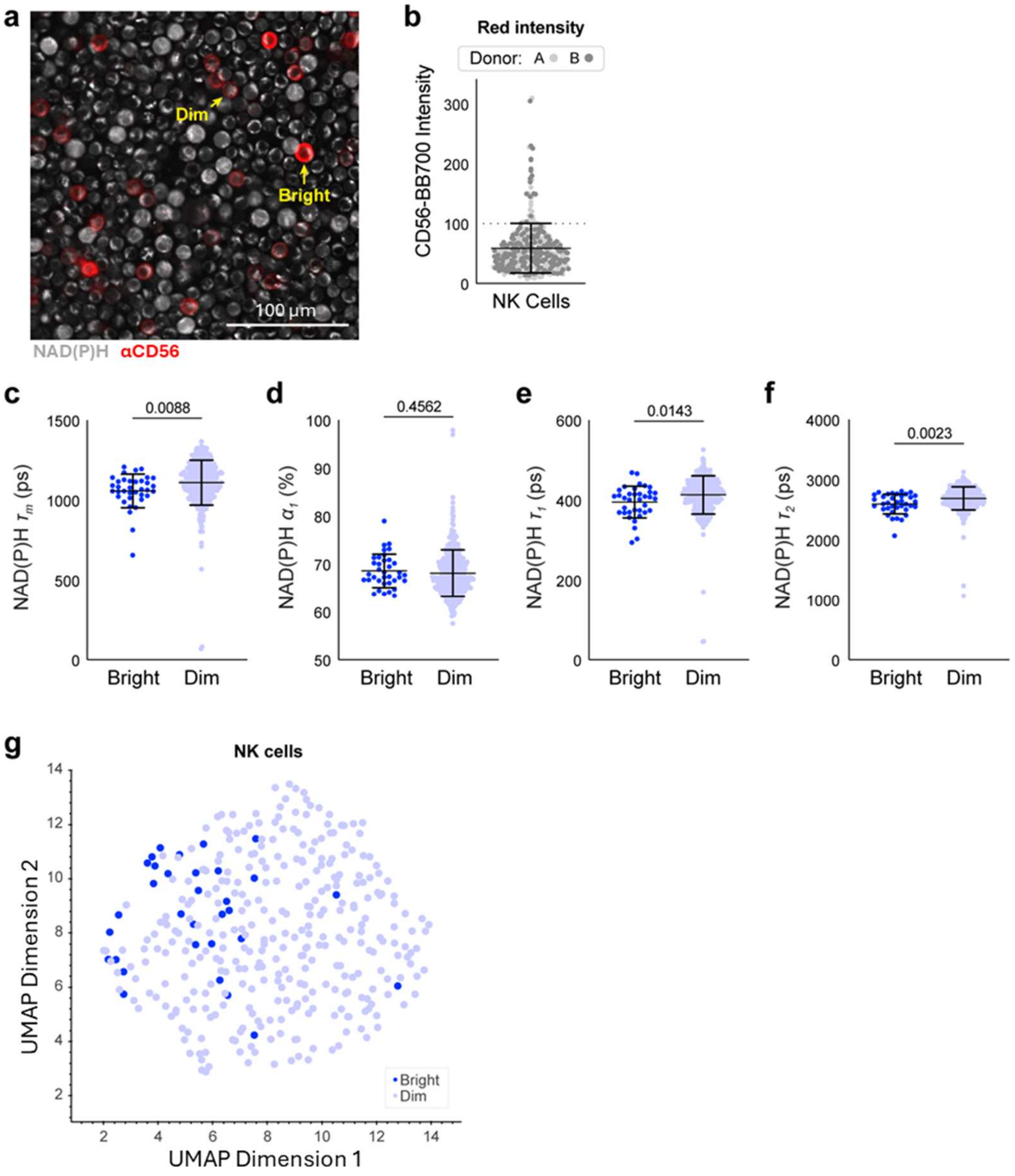


**Supplemental Figure 7. Metabolic subtypes within the quiescent NK cell population.** (a) Representative αCD56 staining (red) overlaid on NAD(P)H intensity (grayscale). (b) An αCD56 intensity value at 100 was selected such that approximately the top 10% brightest CD56-NK cells fell above the threshold. (c-f) NAD(P)H lifetime values showed some significant differences using a two-tailed unpaired t-test. Data given as mean and standard deviation overlaid on single-cell plots. (g) UMAP shows clustering of CD56 bright and dim NK cells using all OMI variables (NAD(P)H and FAD $\tau_m$, $\tau_1$, $\tau_2$, and $\alpha_1$, the ORR, and cell area). n = 327 quiescent NK cells.

| *Target* | *Isotype* | *Clone* | *Reactivity* | *Fluorophore* | *Catalog #* | *Brand* | *Lot* | *Concentration* |
|---|---|---|---|---|---|---|---|---|
| CD4 | Mouse IgG1, κ | SK3 | Human | PerCP-Cyanine5.5 | 344607 | Biolegend | B380254 | 100 µg/mL |
| CD8 | Mouse IgG1, κ | SK1 | Human | PerCP-Cyanine5.5 | 344709 | Biolegend | B356155 | 50 µg/mL |
| CD14 | Mouse IgG2a, κ | M5E2 | Human | PerCP | 301847 | Biolegend | B415977 | 400 µg/mL |
| CD19 | Mouse IgG1, κ | LT19 | Human | Star Bright Blue 700 | MCA1940SBB700 | Bio-Rad | 100004436 | 1.0 mg/ml |
| CD56 | Mouse IgG2b, κ | NCAM 16 | Human | Brilliant Blue 700 | 566574 | BD Horizon | 3226480 | 125µg/mL |
| CD69 | Mouse IgG1, κ | FN50 | Human | PerCP | 310927 | Biolegend | B290414 | 200 µg/mL |

**Supplemental Table 1.** Details of primary antibodies including the host species, clone, manufacturer, and catalog numbers.

| *Cell type* | *Donor* | *Quiescent* | *Activated* |
|---|---|---|---|
| Helper T cells | A | 188 | 78 |
| Helper T cells | B | 161 | 153 |
| Helper T cells | C | 204 | 132 |
| Cytotoxic T cells | A | 160 | 46 |
| Cytotoxic T cells | B | 257 | 135 |
| Cytotoxic T cells | C | 137 | 172 |
| Monocytes | A | 159 | 95 |
| Monocytes | B | 150 | 82 |
| Monocytes | C | 116 | 128 |
| B cells | A | 115 | 110 |
| B cells | B | 116 | 66 |
| B cells | C | 137 | 164 |
| NK cells | A | 170 | 129 |
| NK cells | B | 157 | 18 |
| NK cells | C | 183 | 120 |

**Supplemental Table 2.** Cell counts by cell type, activation, and donor across all experiments.
